# Recurrent neural networks can explain flexible trading of speed and accuracy in biological vision

**DOI:** 10.1101/677237

**Authors:** Courtney J Spoerer, Tim C Kietzmann, Johannes Mehrer, Ian Charest, Nikolaus Kriegeskorte

## Abstract

Deep feedforward neural network models of vision dominate in both computational neuroscience and engineering. The primate visual system, by contrast, contains abundant recurrent connections. Recurrent signal flow enables recycling of limited computational resources over time, and so might boost the performance of a physically finite brain or model. Here we show: (1) Recurrent convolutional neural network models outperform feedforward convolutional models matched in their number of parameters in large-scale visual recognition tasks on natural images. (2) Setting a confidence threshold, at which recurrent computations terminate and a decision is made, enables flexible trading of speed for accuracy. At a given confidence threshold, the model expends more time and energy on images that are harder to recognise, without requiring additional parameters for deeper computations. (3) The recurrent model’s reaction time for an image predicts the human reaction time for the same image better than several parameter-matched and state-of-the-art feedforward models. (4) Across confidence thresholds, the recurrent model emulates the behaviour of feedforward control models in that it achieves the same accuracy at approximately the same computational cost (mean number of floating-point operations). However, the recurrent model can be run longer (higher confidence threshold) and then outperforms parameter-matched feedforward comparison models. These results suggest that recurrent connectivity, a hallmark of biological visual systems, may be essential for understanding the accuracy, flexibility, and dynamics of human visual recognition.

**Author summary:** Deep neural networks provide the best current models of biological vision and achieve the highest performance in computer vision. Inspired by the primate brain, these models transform the image signals through a sequence of stages, leading to recognition. Unlike brains in which outputs of a given computation are fed back into the same computation, these models do not process signals recurrently. The ability to recycle limited neural resources by processing information recurrently could explain the accuracy and flexibility of biological visual systems, which computer vision systems cannot yet match. Here we report that recurrent processing can improve recognition performance compared to similarly complex feedforward networks. Recurrent processing also enabled models to behave more flexibly and trade off speed for accuracy. Like humans, the recurrent network models can compute longer when an object is hard to recognise, which boosts their accuracy. The model’s recognition times predicted human recognition times for the same images. The performance and flexibility of recurrent neural network models illustrates that modeling biological vision can help us improve computer vision.

## Introduction

Neural network models of biological vision have a long history [1–3]. The recent success of deep neural networks in computer vision has led to a renewed interest in neural network models within neuroscience [4–6]. Contemporary deep neural networks not only perform better in computer-vision tasks, but also provide better predictions of neural and behavioural data than previous, shallower models [7–11]. The dominant model class in both computer vision and visual neuroscience is the feedforward convolutional neural network (fCNN).

Inspired by the primate brain, fCNNs employ a deep hierarchy of linear-nonlinear filters with local receptive fields. However, they differ qualitatively from their biological counterparts in terms of their connectivity. Notably they lack the abundant recurrent connectivity that characterises the primate visual system. In terms of recognition behaviour, fCNNs and primates do show similar patterns of image classifications at the level of object categories, but their behaviour diverges when the comparison is made at the level of individual images [12]. Moreover, it has been shown that fCNNs heavily rely on texture in image classification, whereas humans more strongly rely on larger-scale shape information [13].

The initial computations supporting rapid recognition in primates can be modeled as a feedforward process [14]. However, neuroanatomical studies have shown that the primate visual system has a highly recurrent connectivity [15–17]. Recordings of neuronal activity further indicate that the recurrent connections are utilised during object recognition [18–25].

Motivated by the neuroanatomical and neurophysiological evidence, recent modeling work has focused on introducing recurrence into the framework of convolutional neural networks. Recurrent neural networks naturally lend themselves to the processing of temporal sequences, such as dynamic visual sensations. However, even for recognition of static images, recurrent convolutional neural networks (rCNNs) have been shown to bring advantages [26–30]. Recurrence brings performance benefits in object recognition tasks, with recurrent networks outperforming feedforward networks of similar complexity (typically measured by the number of parameters) [26–29]. Moreover, rCNNs are better able to explain neural and behavioural data than their feedforward counterparts [24, 25, 29, 31, 32]. However, performance gains associated with recurrent architectures have previously been shown only for small-scale visual tasks [26–28] or using specialised forms of recurrence [29]. Here we investigated whether rCNNs can outperform feedforward control models matched in their number of parameters on large-scale recognition tasks and on predictions of human reaction times.

Beyond the number of parameters, we must consider the computational cost of recognition. A recurrent network might outperform a feedforward network with a similar number of parameters, but require more cycles of computation and more time to arrive at an accurate answer. Primate brains employ a flexible mechanism that can take more or less time and energy for computations, depending on the difficulty of recognition. This aligns with computational theories of perceptual decision making in primates, which posit that evidence is accumulated until a threshold is reached before making a decision [33]. For some images, fast feedforward computations may be sufficient [24]. If the network converges on a decision in the initial feedforward sweep, then recurrent computation might not be required. For more difficult images, recurrent computation might be essential to ensure accurate recognition. Threshold-based decision making might allow an rCNN to save time and energy on average by only running for the number of time steps required for a given level of confidence.

Threshold-based decision making enables the flexibility of a speed-accuracy trade-off (SAT), explaining an important feature of biological object recognition [34]. A recurrent network can run until it reaches a predefined level of confidence, with the threshold set lower if there is time pressure. The reaction time, then, will reflect both the time pressure (which depends on the situation) and the difficulty of recognition (which depends on the image).

In engineering, a speed-accuracy trade-off might alternatively be implemented using a range of separate neural network models of varying scale [35, 36]. However, using multiple models to implement an SAT has three disadvantages: (1) It requires more storage. (2) It requires the selection of the appropriate model for each scenario at the start of the process. (3) Once the model is chosen the reaction time is fixed and the model cannot flexibly choose to compute longer for harder images. Threshold-based decisions, thus, appear advantageous for both biological and artificial vision, which similarly face limitations of space, time, and energy.

To better understand the role of recurrent computations, we compared rCNNs to feedforward (fCNN) control models in terms of their object-recognition performance and their ability to account for human visual recognition behaviour. We trained our networks on the ImageNet Large Scale Visual Recognition Challenge (referred to as *ImageNet* for brevity) [37], and a more ecologically valid recognition task called *ecoset* [38]. We investigated whether recurrence improves recognition accuracy in these tasks. We further modelled threshold-based decision making in the rCNNs, varying the threshold to control the SAT [34], and compared reaction times to different images between rCNNs and human observers.

## Results

We trained a range of deep convolutional neural networks on two large-scale visual object-recognition tasks, ImageNet [37] and ecoset [38]. The networks trained included a feedforward network, referred to as B (for *bottom-up* only), and a recurrent network, referred to as BL, with *bottom-up and lateral* recurrent connections (recurrent connections within a layer). We focus our investigation on lateral connections, which constitute a form of recurrence that is ubiquitous in biological visual systems and proved powerful on simple tasks in our earlier work [28].

The rCNNs were implemented by unrolling their computational graphs for a finite number of time steps (see *Methods*). Each model was trained to produce a readout at each time step, which predicts the category of the object present in the image.

Adding recurrent connections to a feedforward model increases the number of parameters. We therefore used three larger feedforward architectures that were approximately matched in the number of parameters (Fig. 1) as control models. Control models were matched in the number of parameters by increasing (1) the size of the convolutional *kernels*, (2) the number of *feature maps*, and (3) the *depth* of the network (referred to as B-K, B-F and B-D, respectively, where the B indicates that these models had only *bottom-up* connections). Parameter matching is important because parameters are costly. Both engineering and biology must consider two main costs that scale with the number of parameters: the space requirements for storing the parameters and the data requirements for setting the parameters.

**Fig 1.**
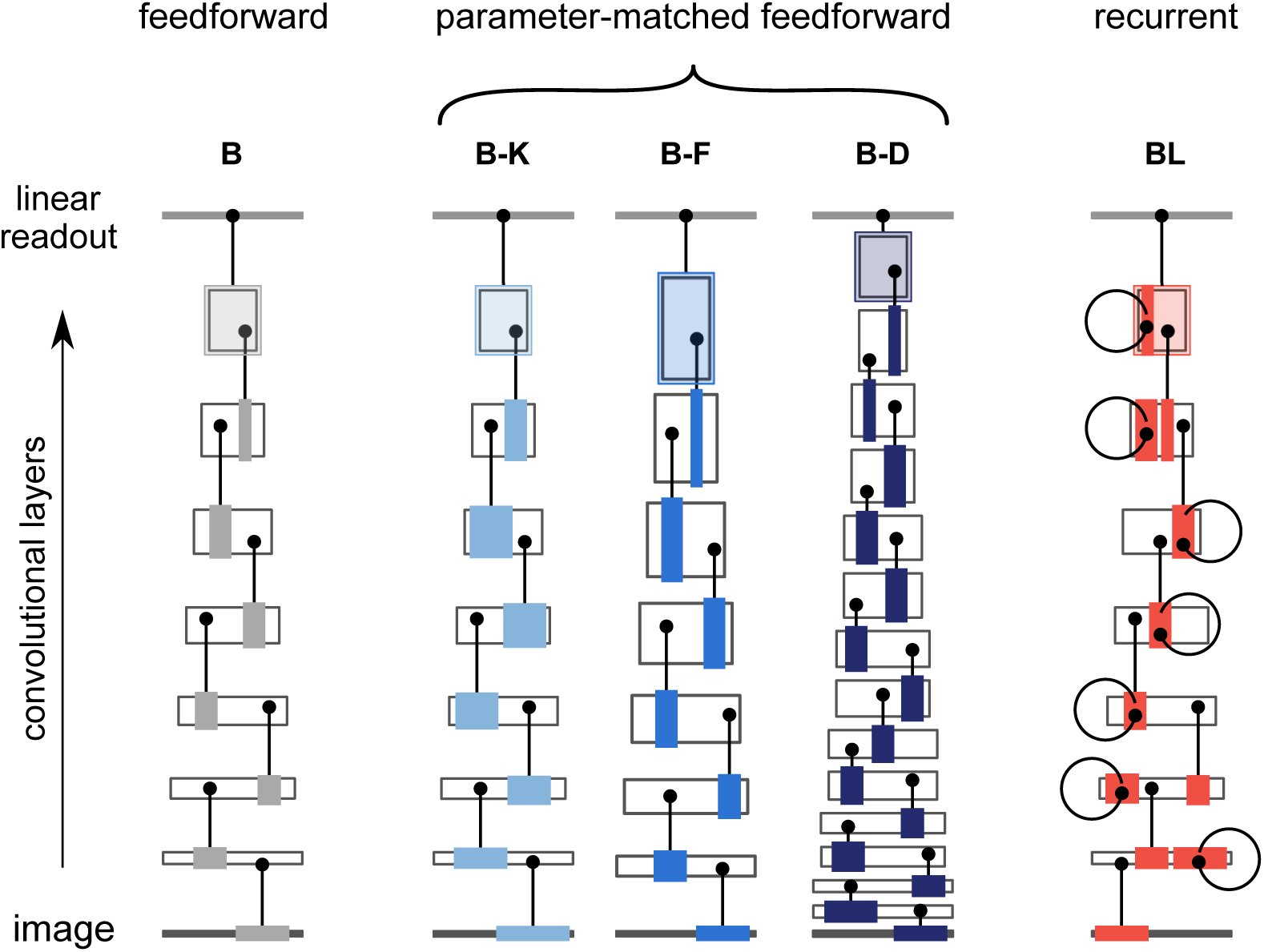
Schematic representation of the parameter-matched networks. White boxes represent convolutional layers, with the width representing the spatial dimensions of the convolutional layers and the height representing the number of feature maps. Models were matched in the number of parameters by increasing (1) the size of the convolutional *kernels* (B-K), (2) the number of *feature maps* (B-F), and (3) the *depth* of the network (B-D). Example units (black dots) are linked to coloured regions representing their input kernels (which differ in width in B-K). The extents are illustrative and not drawn to scale.

A major benefit of recurrent models is that they can run more computations without requiring more parameters. The computational graph of a recurrent model grows with the number of time steps the model runs for. The total number of computations (whether performed in parallel or sequentially) and the maximum number of sequential nonlinear transformations (which we refer to as the *computational depth*), therefore, are limited by the number of time steps, not by the number of layers, in a recurrent model. However, a feedforward architecture can also achieve any prespecified number of computations and computational depth by including enough units and layers. This raises the question of how a feedforward model with a matched computational graph compares to an rCNN. We therefore trained a further feedforward control model whose architecture was defined by unrolling the rCNN. This model (referred to as B-U, for *bottom-up unrolled*) has an identical computational graph (and thus the same number of computations and computational depth), but unique parameters for each convolution (i.e. no weight sharing across time). As a result, B-U has more than seven times as many parameters as BL (212.7 million for B-U, 28.9 million for BL). B-U was trained with category readouts at regular intervals throughout the network (matching the readouts at the end of each time step in BL). Including multiple readouts allows B-U to explain variability in human reaction times by terminating at different stages.

It is possible to alter the number of parameters and computations in the networks by including other architectural features such as adding Inception modules [39]. However, to ensure a meaningful comparison, we aimed to maintain as close a similarity as possible between recurrent and feedforward architectures. The pros and cons of the different control models are outlined in Table 1 (see *Methods* for a detailed description of the models and training procedures).

**Table 1.** Pros and cons of different control models

| **Control model matched in *number of parameters*** |  |
| --- | --- |
| Larger *kernels* (B-K) | **Pro:** Matches the number of units in each layer and the number of layers.<br>**Con:** Inefficient use of parameters in relation to object recognition performance. |
| More *feature maps* (B-F) | **Pro:** Matches the number of layers and better performance gains than increasing kernel size.<br>**Con:** Does not match the number of feature maps in each layer and has worse object recognition performance than making the network deeper. |
| Greater *depth* (B-D) | **Pro:** Tends to yield best improvement in performance for additional parameters.<br>**Con:** Does not match the number of layers in the recurrent model. |
| **Control model matched in *computational graph*** |  |
| Feedforward network matching the *unrolled* recurrent network (B-U) | **Pro:** Matches the computational graph and thus, in particular, the number of computations and the computational depth.<br>**Con:** The number of parameters grows precipitously with the number of time steps of the recurrent model, and ends up being much larger than in the recurrent model. |

### Recurrent networks outperform parameter-matched feedforward models

We compared the performance of the recurrent BL architecture to the baseline feedforward, and parameter-matched control architectures. For each architecture, we trained and tested separate models on the ImageNet and ecoset visual recognition tasks. For the recurrent BL networks, we defined the prediction of the model as the average of the category readout across all time steps, which we refer to as the *cumulative readout*. The cumulative readout tends to produce superior performance (see *Methods*). Top-1 accuracies are used throughout.

The recurrent models outperformed the baseline and all parameter-matched feedforward models (Fig. 2B). BL showed a performance benefit of about 1.5 percentage points relative to the best parameter-matched feedforward model, B-D, on both datasets (Table 2).

**Table 2.** Accuracies on held-out data for parameter-matched models

| models | ImageNet | ecoset | parameters |
| --- | --- | --- | --- |
| B (baseline) | 58.42% | 64.25% | 11.0 million |
| B-K (larger kernels) | 56.46% | 62.81% | 39.8 million |
| B-F (more feature maps) | 60.34% | 66.54% | 40.0 million |
| B-D (deeper network) | 62.68% | 68.36% | 28.9 million |
| BL (recurrent) | **64.37%** | **69.98%** | 28.9 million |

**Fig 2.**
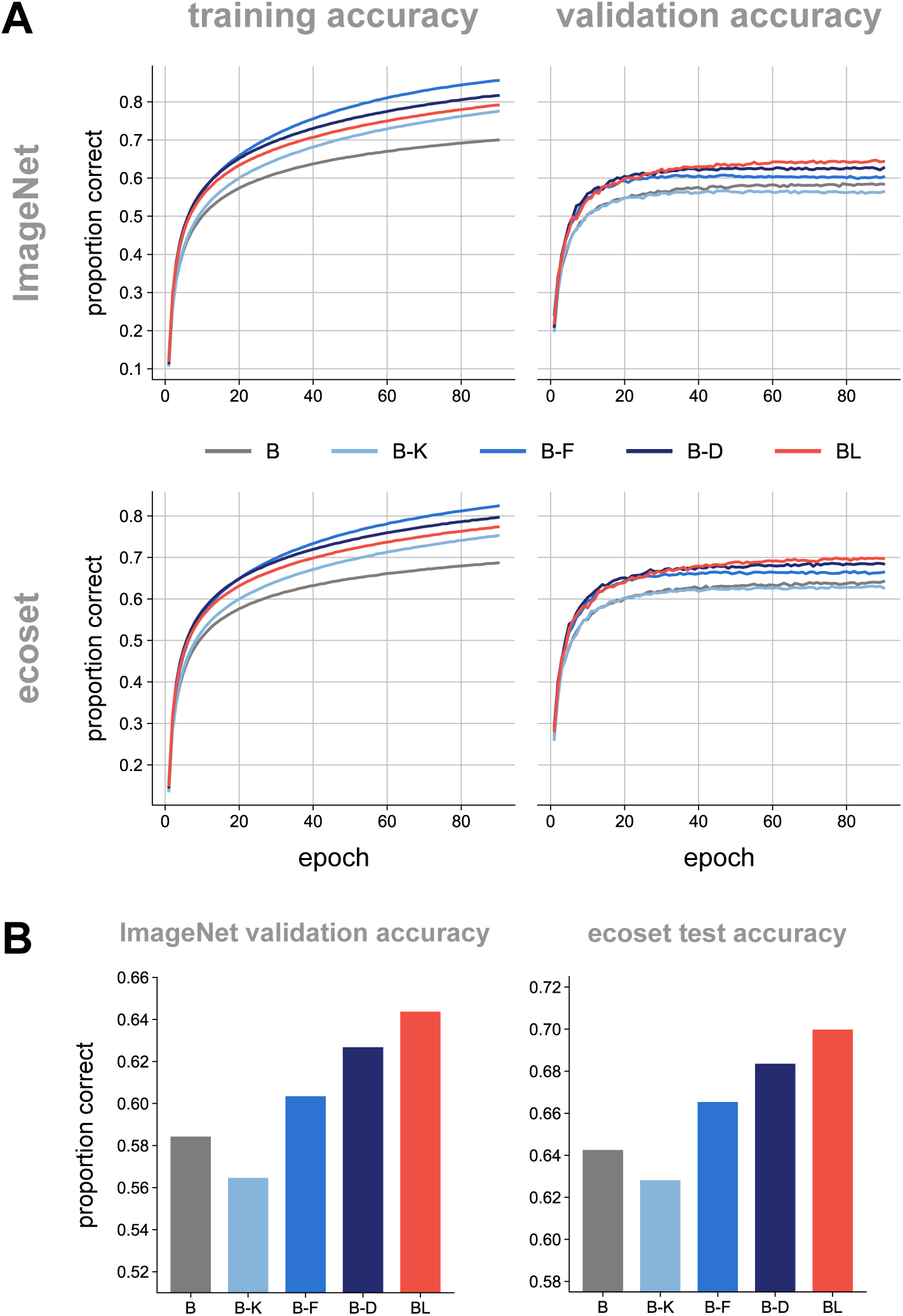
ImageNet and ecoset task performance for rCNN and parameter-matched controls. Our rCNN model (red) achieves higher validation accuracy than parameter-matched control models (shades of blue). (A) Training and validation accuracies across training epochs for all networks (top-1). (B) Performance of networks on held-out data using the fully-trained networks. All pairwise differences in model performance were significant (*p ≤* 0.05, McNemar test, Bonferroni corrected for all pairwise comparisons).

The number of parameters are calculated for ImageNet models, ecoset models have slightly fewer parameters due to fewer categories in the final readout layer.

Both B-D (deeper network) and B-F (more feature maps) outperformed the baseline model, B. B-K has a worse test accuracy than the baseline model but a higher training accuracy (Fig. 2A). This suggests that using additional parameters to increase the kernel size in our models leads to overfitting rather than a generalisable increase in performance.

Pairwise McNemar tests [40, 41] showed all differences in model performance to be significant (*p ≤* 0.05, corrected). Bonferroni correction was used to correct for multiple comparisons in order to control the family-wise error rate at less than or equal to 0.05.

### A recurrent model with entropy thresholding predicts a speed-accuracy trade-off

Across recurrent computations in our rCNNs, the probability mass of the output distribution tends to concentrate, indicating that the network’s confidence in its classification is rising. We used the entropy of the output distribution to measure the network’s confidence. Zero entropy would indicate that the network is certain, with all probability mass concentrated on a single class. The network runs until the entropy of its cumulative readout falls below a predefined entropy threshold. The final cumulative readout is then taken as the network’s classification.

Entropy thresholding has the benefit of being economical, as it uses the minimum number of time steps to reach the required level of confidence for an image. Moreover, entropy thresholding is related to neuroscientific theories of decision making, where evidence is accumulated until it reaches a bound [33].

At a given entropy threshold, a recurrent model may choose to compute longer for harder images. The model’s reaction time (i.e. the number of time steps required to reach the entropy threshold) thus varies across images. For a given rCNN, the reaction time is proportional to the computational cost of recognising an image (i.e. the number of floating-point operations), and thus to the energy cost, which might be related to the metabolic cost in a biological neural network.

For each setting of the entropy threshold, we estimated the accuracy and the computational cost. We estimated the accuracy as the overall test-set accuracy at this threshold. We estimated the expected computational cost as the average, across the test set, of the number of floating-point operations used. We plotted the accuracy of the model as a function of the computational cost (Fig. 3). For a given recurrent model, the resulting plot reflects a speed-accuracy trade-off, because the reaction time is proportional to the computational cost. Across thresholds, the accuracy rises with the average time taken (and average computational cost), until it saturates.

**Fig 3.**
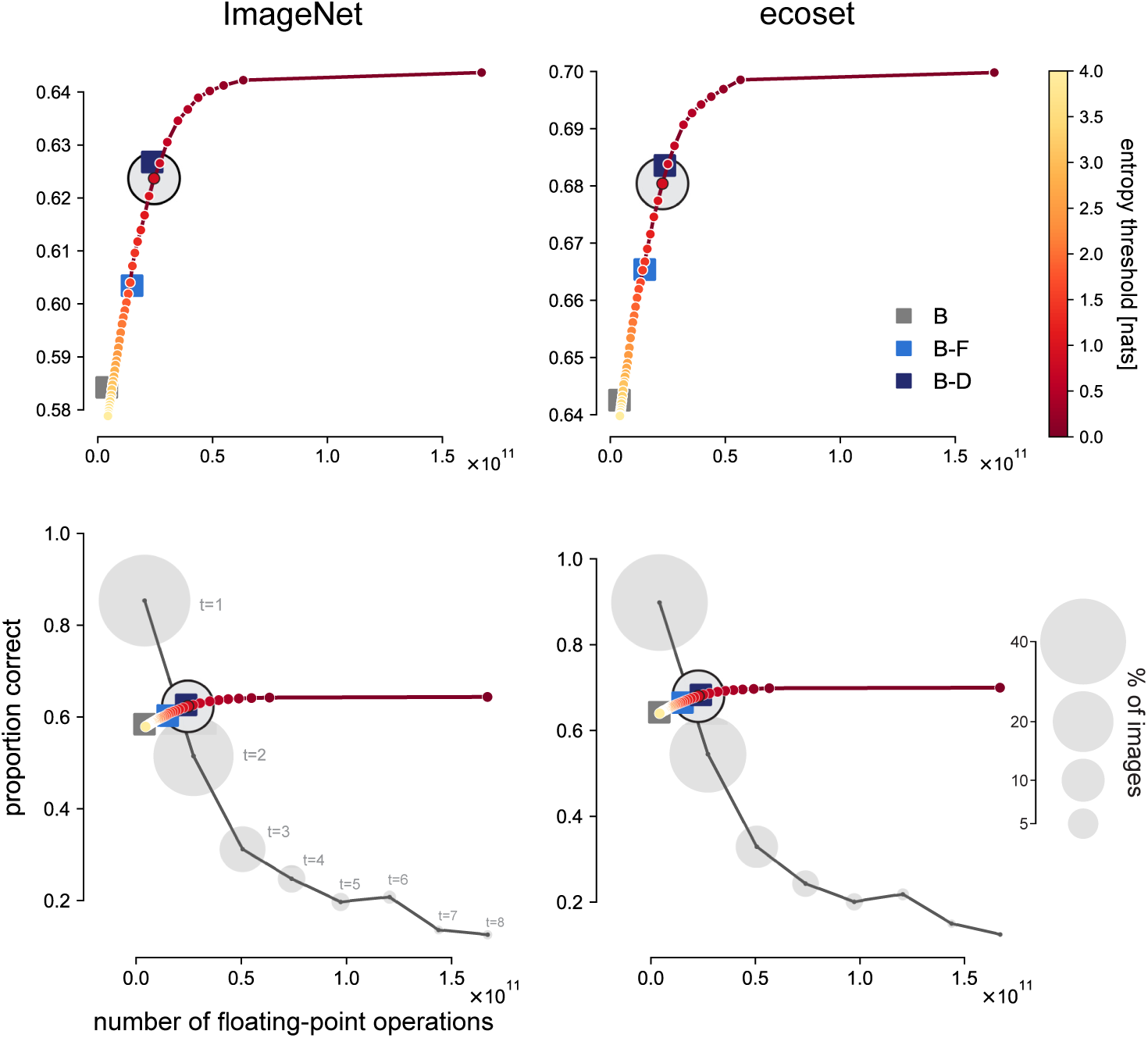
Validation accuracy as a function of computational cost for feedforward and recurrent models. Each feedforward model (squares in shades of blue) requires a fixed number of floating-point operations for a single sweep of computation. The top row shows that feedforward models requiring more computation (horizontal axes) had higher top-1 validation accuracy (vertical axes). The recurrent models (yellow-to-red line) could be set to terminate at different levels of confidence, specified as the entropy of the softmax output. For each entropy threshold (colour bar), the computational cost (mean number of floating-point operations) and the top-1 validation accuracy (proportion correct) were computed across the test set. The recurrent models could flexibly trade speed for accuracy (lines in top panels). They achieved the same accuracy as each feedforward control model when given a matched computational budget, and greater accuracy than any of the feedforward models when run longer. The bottom panels replot the data shown in the top panels and additionally show, for a single entropy threshold of the recurrent models, how computational cost varies across images (horizontal domain of the black lines) and what accuracy is achieved at each computational cost. The black line shows the accuracy as a function of computational cost for the selected entropy threshold. The area of each gray circle is proportional to the percentage of images for which the model reaches the entropy threshold at a given computational cost. The open black circle is the average of the points on the black line, weighted by the percentage of images for each computational cost. We see that, at the selected entropy threshold, the model responds rapidly for about half of the images and achieves high performance on these “easy” images. It computes longer for “hard” images, balancing the cost of lower accuracy against the cost of greater expenditure of energy and time.

**Fig 4.**
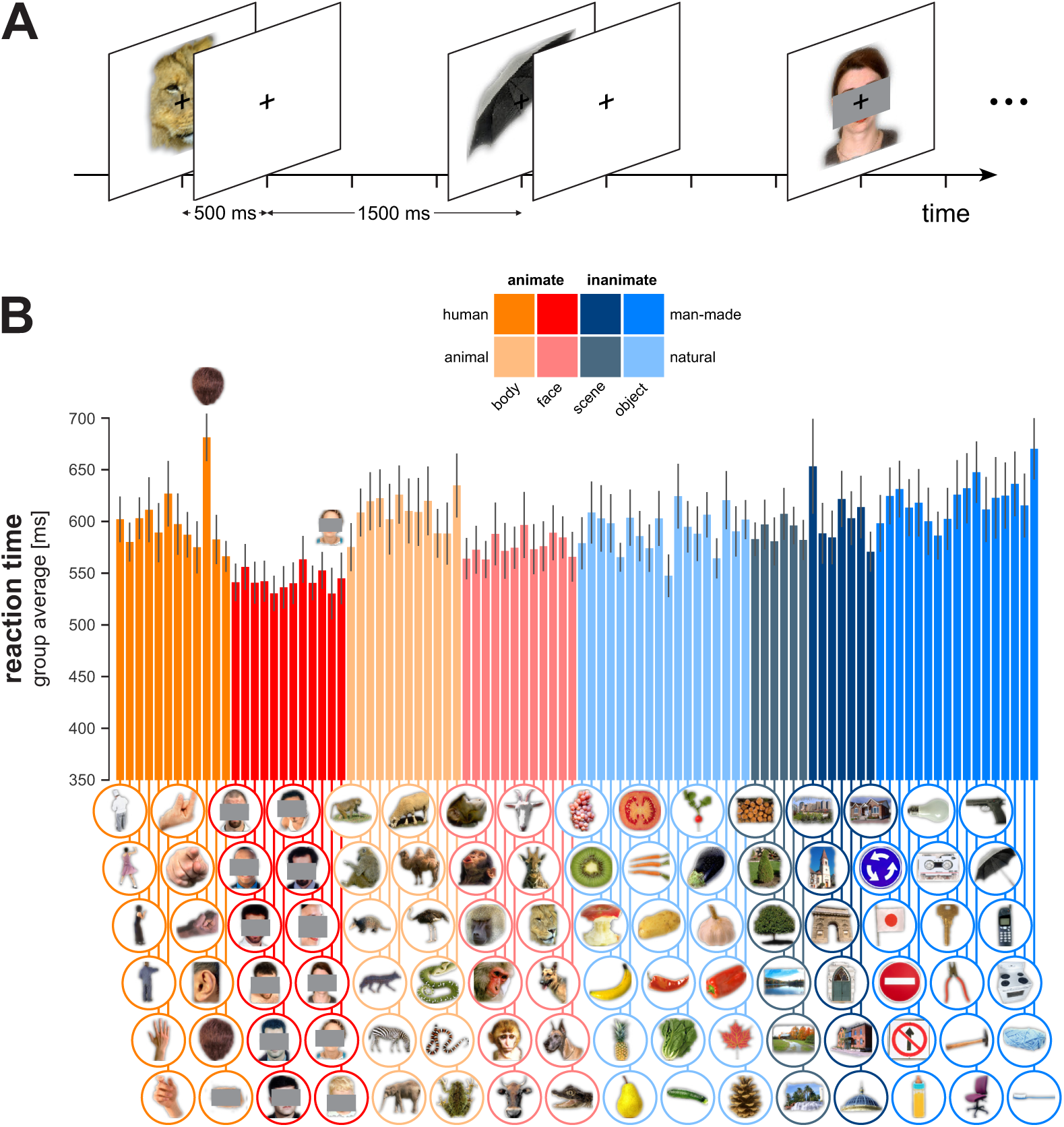
Human behavioural experiment. (A) Human subjects were presented with images of isolated objects of different categories and classified the images as animate or inanimate by pressing one of two buttons on each trial. (B) Group-average reaction time for each image. Error bars show the standard error of the mean. [Human faces are partially occluded in this figure to comply with the requirements of bioRxiv, but were presented to human subjects and models without the gray occluder boxes.]

### A single rCNN emulates the accuracies of different fCNNs when its confidence threshold is set to match the fCNN’s computational cost on average

We also assessed the accuracy and computational cost of the feedforward models. Results are shown in the context of those for the recurrent models in Fig. 3. Feedforward models are represented by single points because their computational cost is constant.

When comparing the recurrent models to the feedforward models, we see a remarkable correspondence between the two classes of architecture: The points describing the feedforward models fall on the line describing how the recurrent model trades off speed and accuracy: Given the computational budget of a particular feedforward model, the recurrent model achieves the same accuracy. However, the computational costs and accuracies of the feedforward models are fixed, whereas recurrent models can be left to compute longer. Given a larger computational budget, the recurrent model will achieve higher accuracy than any of the parameter-matched feedforward models.

To inferentially compare the performance of the feedforward and recurrent networks at matched computational cost, we considered the performance of the recurrent networks at a single entropy threshold. We selected the threshold that minimises the absolute difference between the average number of operations for the recurrent network and the number of operations for the feedforward network. McNemar tests were again used to compare the performance of the networks.

Across both datasets only one significant difference in performance was found between recurrent and feedforward models. This difference was the between B and BL in ImageNet, which achieved 58.42% and 57.71%, respectively, a difference of 0.70% (*p <* 0.001, uncorrected). This comparison matches a pass through B to the initial feedforward pass through BL. BL appears to slightly compromise its performance on the initial feedforward pass to support later gains through recurrence. All other differences between BL and feedforward networks were even smaller and not significant, ranging between −0.37% and +0.32%, relative to the performance of BL. B-K was excluded from this analysis because it had worse performance than the baseline feedforward model (possibly due to overfitting).

These results suggest that recurrent models perform similarly to feedforward models when allowed the same average number of floating-point operations. This may be surprising given that recurrent models must use the same weights across multiple time steps, whereas feedforward models do not face this constraint. We may have expected the operations learned by recurrent models to be less efficient with regard to performance achieved at a given computational cost. Instead, we found that the computational efficiency of recurrent and feedforward networks are well matched. The graceful degradation of performance of recurrent models when the computational cost is limited may depend on training with a loss function that rewards rapid convergence to an accurate output (see Methods). Recurrent models may benefit from the fact that they can save computation on easy images, enabling them to expend greater computational cost than their feedforward competitors on harder images, while matching the average computational cost.

Overall our results suggest that we can use a single recurrent network to flexibly emulate the accuracies achieved by different feedforward models. Matching the accuracy of a given feedforward model will come at a computational cost that approximately matches the computational cost of the feedforward model on average. The recurrent model will terminate faster for easy images and compute longer for harder images. The recurrent model can also be set to run more recurrent computations enabling it to achieve higher performance than the parameter-matched feedforward networks.

### Reaction times from recurrent networks better explain human reaction times

Recurrent connections endow a model with temporal dynamics. If the recurrent computations in a model resemble those of the human brain at some level of abstraction, then model behaviour should be predictive of human behaviour. For example, images that take longer for the model to recognise should also take longer for humans to recognise.

To test this hypothesis we used data from an object categorisation task where humans had to categorise 96 full-colour images as animate or inanimate. Reaction times were recorded from 20 human participants. Our goal was to quantify the extent to which model reaction times predicted human reaction times.

We fitted recurrent and feedforward models to these human data and tested the fitted models using cross-validation across images and subjects. Feedforward models were included in this analysis to test the competing hypothesis that varying reaction times could be explained by halting computations part way through the feedforward sweep. The feedforward models tested included a deep feedforward control model matched to BL in terms of the computational graph (B-U). B-U is identical to a BL network unrolled across time, except for the fact that it is not constrained to recycle its parameters across time steps. The B-U model had category readouts at intermediate layers, matching BL’s readouts at multiple time steps. Additional feedforward models were also used including, B-D (trained on ImageNet and ecoset) and feedforward models pre-trained on ImageNet that are popular in the machine learning literature [42–47].

The models were fitted to the human data in two stages: (1) An animacy discrimination readout was fitted. (2) An entropy threshold was fitted to enable measurement of model reaction times. To fit the animacy discrimination readout, eight readouts were placed at regular intervals throughout the networks. The readouts were trained to maximise performance on the animacy discrimination task using a separate set of images from those used in the human behavioural task. The entropy threshold was fitted to maximise the Pearson correlation between network and human reaction times. We used a double leave-one-out cross-validation approach, ensuring that thresholds were fitted using data from one set of images and subjects, and model reaction times compared to human reaction times for an independent set of images and subjects. The network reaction time was taken as the position of the readout that first reached the entropy threshold. This procedure resulted in a predicted reaction time for each subject-image pair.

To compare the ability of different models to predict human reaction times, we computed the correlation between network reaction times and the reaction times for individual subjects. A human consistency metric was also computed by correlating the reaction times of a single human participant against the average of all other human participants. This procedure provides a lower bound on the noise ceiling, i.e., a lower bound on the performance that the true model would achieve given the noise and intersubject variability [48]. Correlations between model and human reaction times, as well as human consistency (lower bound of the noise ceiling), are shown in Fig. 5.

**Fig 5.**
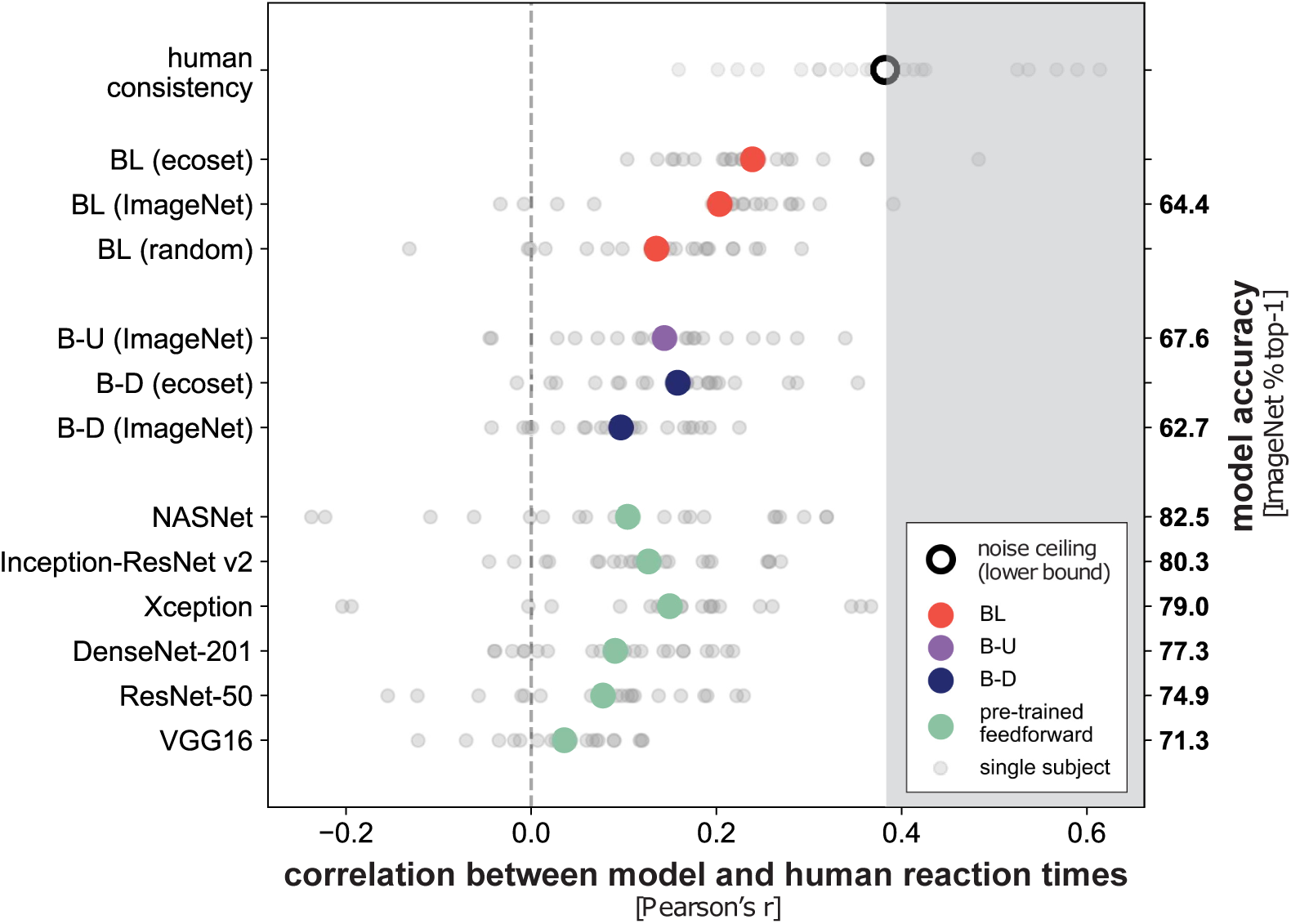
Reaction times from recurrent networks explain human reaction times better than feedforward networks. Small grey dots represent the Pearson correlation between the network and single subject reaction times. Large dots represent the mean correlation across subjects. Human consistency (black circle) provides a lower bound on the noise ceiling and is computed by correlating reaction times for a single subject with the average reaction time for all other subjects. For each network, multiple sigmoid animacy readouts were placed at even intervals throughout the networks. Animacy readouts were trained to maximise accuracy using a separate set of images not used in the human behavioural experiments. For each model, an entropy threshold was fitted, using independent subjects and images, so that model reaction times best predicted human reaction times (cross-validation).

Paired two-tailed permutation tests were used to detect significant differences in reaction time correlations between networks. The Benjamini-Hochberg procedure was used to account for multiple comparisons by controlling the false discovery rate at 0.05 [49].

The results show that reaction times extracted from BL trained on ecoset best predicted human reaction times, outperforming all feedforward networks and the untrained BL network (FDR *q <* 0.05). Notably, the explanatory benefit over the feedforward architectures includes the control model B-U, which is highly similar to BL, but requires the training and storage of a significantly larger number of parameters (212.7 million for B-U compared to 28.9 million for BL, Fig. 6). While this significantly larger model, perhaps not surprisingly, yields better overall task performance, it is outperformed by BL in its ability to mirror human reaction times.

**Fig 6.**
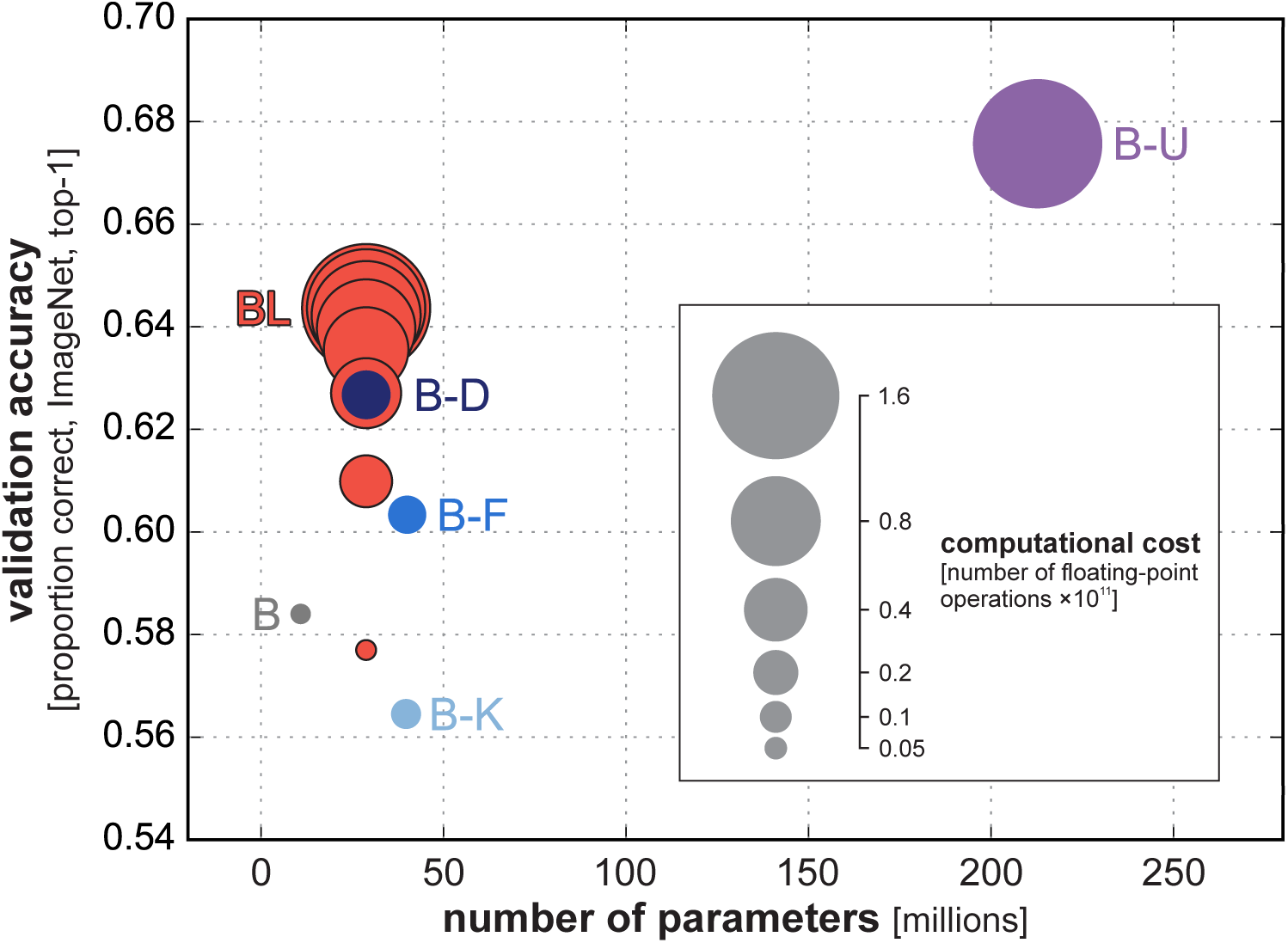
Relationship between validation accuracy, number of parameters and computational cost across models. The validation accuracy (vertical axis) is the proportion top-1 correct classifications of the trained models on ImageNet. For each model (coloured disc), the validation accuracy is plotted against the number of parameters (horizontal axis). The area of the coloured discs is proportional to the computational cost as measured by the number of floating point operations required to run the model. The red circles correspond to different numbers of recurrent cycles of computation of the BL recurrent convolutional network. For model abbreviations (B, B-K, B-F, B-D), see Fig. 1. B-U is the unrolled control model, with a computational graph matched to BL, but no parameter sharing across cycles of computation.

BL trained on ImageNet predicted the human reaction times better than all feedforward networks (*p <* 0.05, FDR corrected) apart from Xception and B-D trained on ecoset, where there was no significant difference. Relative to the randomly initialised BL model, all feedforward models were either significantly worse at explaining human reaction times or there was no significant difference in correlation (FDR *q <* 0.05). B-D trained on ecoset had a significantly higher correlation than B-D trained on ImageNet (FDR *q <* 0.05). All models had a significantly lower correlation that the human consistency metric (FDR *q <* 0.05).

In summary, the comparison of model reaction times to human reaction times demonstrated the benefits of recurrent processing compared to all other networks tested. The recurrent BL model also explained reaction times better than the B-U model, although B-U had the same computational graph and matched readouts at intermediate stages.

### Exploratory analysis of lateral connectivity patterns

To better understand the lateral connectivity patterns that emerge from category training in our recurrent models, we analysed the recurrent connections in the first network layer of a BL network trained on ImageNet. The focus on the lowest network layer enabled us to visualise connectivity patterns in the pixel space. Our goal was to qualitatively assess similarities to intra-area connectivity in primate V1. To summarise the large number of lateral connections in the first network layer alone (over 450,000 connections), we used principal components analysis (PCA), decomposing the lateral-weight templates into orthogonal components (similar to Linsley et al. [30], see *Methods* for details). We then visualised these lateral-weight components together with the bottom-up features that they connect. Fig. 7 shows the first five weight components (capturing 43% of variance across all recurrent weights). Although in-depth confirmatory analyses of the learned connectivity are out of the scope of the current work, it is noteworthy that all five components could be interpreted in terms of biological phenomena: inhibition/excitation (component 1), vertical antagonism (component 2), centre-surround antagonism (component 3), horizontal antagonism (component 4), and perpendicular antagonism (component 5). These features could relate to properties of biological visual systems such as border-ownership [50] and contour integration [51] (for a more detailed description of these results, see S1 Text).

**Fig 7.**
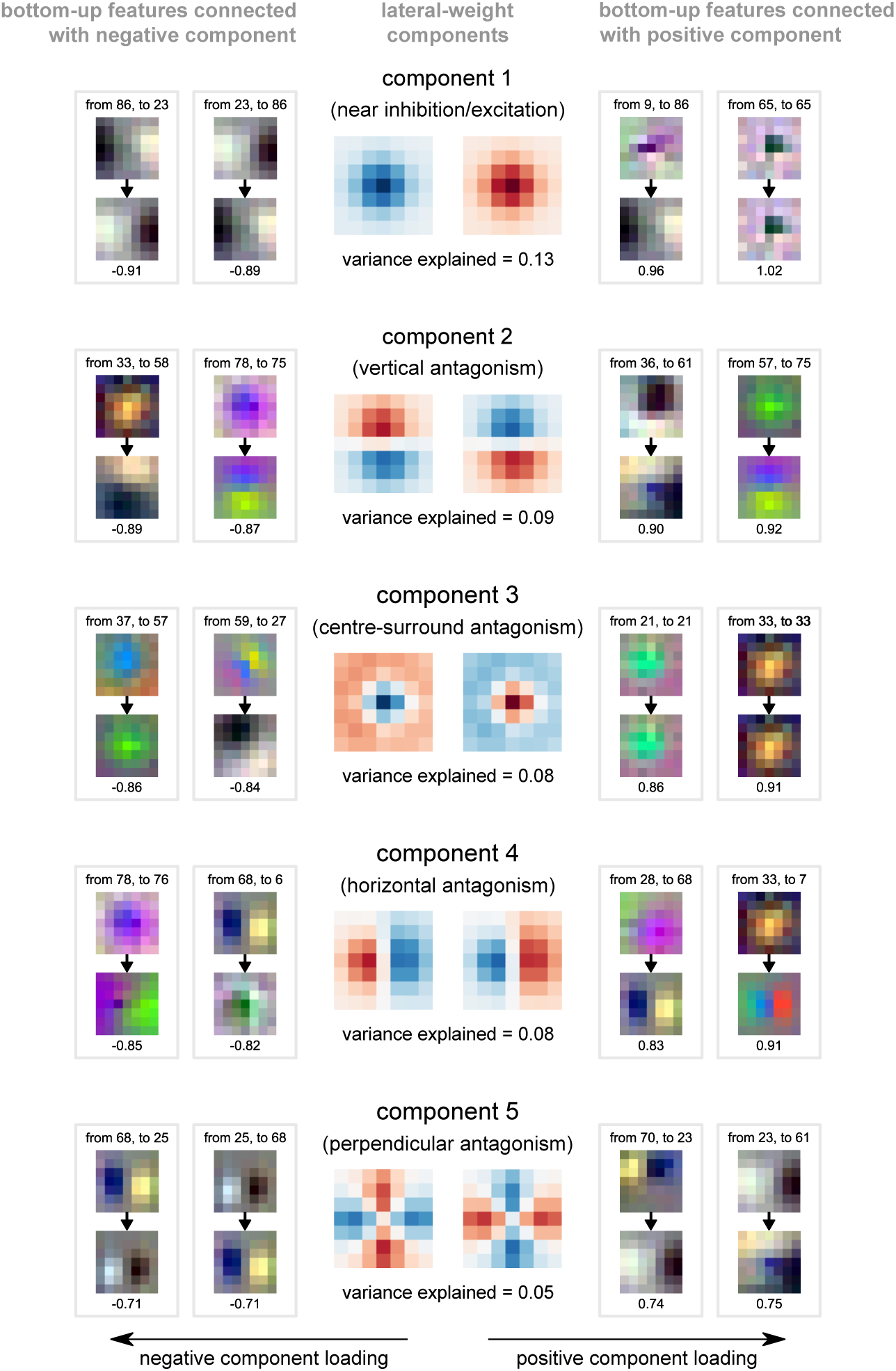
Lateral-weight components for layer 1 of an rCNN trained on ImageNet. Every unit receives lateral input from other units within and across feature maps via a local lateral-weight pattern. We used principal component analysis to summarise the lateral-weight patterns. The top five lateral-weight principal components are shown in both their positive (centre right) and negative forms (centre left). Blue shading corresponds to negative values and red to positive. The proportion of variance explained is given beneath each lateral-weight component. Bottom-up feature maps connected by lateral weights with the strongest positive (right) and negative loadings (left) on the weight component are shown alongside. Arrows between bottom-up features indicate the direction of the connection and the loading is given underneath each pair of bottom-up features.

## Discussion

Our results show that recurrent convolutional neural network models can outperform parameter-matched feedforward convolutional models of similar architecture on large-scale naturalistic visual recognition tasks. In addition to superior performance, rCNNs more closely resemble biological visual systems in both structure and function. Structurally, biological visual systems and rCNNs share ample recurrent signal flow. Functionally, biological visual systems and rCNNs both exhibit greater accuracy and flexibility than fCNNs of similar parametric complexity.

### Flexible trading of speed and accuracy

An important functional feature of our rCNNs is the flexibility to trade off speed and accuracy, which these models share with biological visual systems. The required confidence can be specified in the form of the entropy of the model’s posterior. Recurrent computation is terminated when the posterior probability mass has concentrated such that its entropy dips below the threshold. Recurrent computation will be brief for easy images, for which the model quickly achieves a high-confidence classification. For harder images, recurrent computation can proceed longer.

We expected that the rCNN’s flexibility to read out the category earlier or later would incur a significant cost in terms of accuracy at a given computational budget. Indeed when the number of recurrent cycles of computation is fixed, so as to match the computational cost of a given feedforward network, the accuracy is somewhat lower (Fig. 6, compare dark blue disc for B-D similarly sized red disc for BL). An rCNN trained to flexibly trade speed and accuracy might compromise its performance at a fixed number of time steps, relative to a fCNN with a similar computational budget. However, an rCNN that halts computation when a predefined confidence threshold is reached will terminate early for easy images, saving computation on average. These savings enabled the flexible rCNNs here to achieve the same accuracy as parameter matched fCNNs at the same average computational cost.

We compared the rCNN to a range of fCNNs that had a similar number of parameters, but required a different amount of computation (Fig. 3, top row). The fCNNs requiring more computation achieved higher accuracy. When the rCNN’s confidence threshold was set so as to match any of the fCNN’s accuracy, the average computational cost of the rCNN matched the computational cost of the fCNN.

### Prediction of human reaction times

When a recurrent model is given the ability to trigger perceptual decisions when it reaches a confidence threshold, it will exhibit variable reaction times for different images. This enables recurrent models to make predictions about human reaction times in visual recognition tasks. Feedforward models, by contrast, expend the same amount of time (and computation) for each image, whether easy or hard. As a result, they do not, by default, predict variations in reaction time across images.

It is possible to model categorisation reaction times on the basis of a static representation of the stimuli in a layer of a feedforward neural network. To achieve this, we can assume that the feedforward computation is noisy and therefore must proceed repeatedly while an evidence accumulator averages away the noise. How long the evidence must be accumulated for will depend on where the stimuli fall relative to the decision boundary of our decoder. If a stimulus falls far away from the decision boundary on the correct side in the multivariate response space, then the evidence will accumulate rapidly [33]. We predict fast responses to stimuli far from the decision boundary and slow responses to stimuli close to the boundary [52, 53]. Note, however, that a feedforward model does not inherently require an evidence accumulator. Assuming separate modules for feedforward transformation and evidence accumulation is not well motivated from either a biological or a computational perspective.

Here we went beyond this approach and built recurrent models that naturally produce reactions at different latencies, reflecting the time needed for the posterior probability mass to concentrate on a category such that the posterior entropy reaches a threshold. With this approach, we need not assume an evidence accumulation process external to the model in order to predict reaction times. Moreover, the recurrent inference mechanism is not limited to accumulating a unidimensional noisy evidence signal. Instead, the network can learn more complex recurrent inferential computations as required for the task it is optimised to perform. In our rCNN models here, longer reaction times do not reflect the need to average a weaker noisy signal. Instead, longer reaction times reflect the need for deeper computation on difficult images. The required cycles of recurrent iterative processing delay the response.

In order to be able to compare recurrent and feedforward models, we enhanced the feedforward models by readouts at different stages of feedforward computation. As for the recurrent models, we then predicted the reaction time from the stage at which the posterior entropy hit the threshold. This enabled a fair and direct comparison between recurrent and feedforward models in terms of their ability to predict human reaction times. The recurrent models outperformed the feedforward models at predicting human reaction times (Fig. 5). In particular, the unrolled feedforward model B-U, matched to the BL rCNN in terms of its computational graph, was not able to predict human reaction times as well as the BL rCNN. This suggests that the recurrent use of the same connections, an iterative computation, may be important for explaining human reaction times to particular images.

### Superior accuracy of recurrent models

The performance of recurrent models, relative to feedforward, is consistent with previous work using small-scale machine learning tasks [26, 28]. However, it contrasts with more recent results suggesting that specialised recurrent architectures, in the form of reciprocally gated cells, are required for recurrent networks to outperform their feedforward counterparts in naturalistic visual recognition tasks [29]. One potential explanation of these ostensibly diverging results is the scale of the feedforward control models relative to the recurrent networks. In the experiments described here, the recurrent networks had approximately 72-100% of the parameters of the feedforward control models. In comparison, the baseline recurrent models “Vanilla RNN” (similar to BL) had approximately 39% and 45% of the parameters of the feedforward control models (“FF Deeper” and “FF Wider”, respectively) in [29]. While reciprocally gated cells clearly produce better task performance, this difference in the number of parameters may explain why our recurrent convolutional networks (without the addition of gating) were able to outperform the parameter-matched feedforward models. It also highlights the difficulty of defining appropriate feedforward control models. Here, we took the approach of matching the number of parameters in feedforward and recurrent models. We additionally considered the performance of a fCNN model (B-U) with the same computational graph as the rCNN. The latter approach has the advantage of matching the number of computations and the computational depth, but it has the disadvantage of a severe mismatch in the number of parameters (larger by factor 7 in the fCNN here).

### Biology and engineering

Our rCNN models borrowed two ideas from the literature on biological decision making: threshold-based decision making [33] and speed-accuracy trade-offs [34]. First, using a fixed posterior-entropy threshold, networks were able take longer to recognise more difficult images. Second, by varying the posterior-entropy threshold, networks could change their required confidence, trading off accuracy for speed. These behaviours enable economical object recognition, only spending the time (and energy) required by the given task or situation. The type of flexible behaviour demonstrated here for rCNNs is useful in both biological and artificial object recognition, where time and computational resources for inference are often limited. Vision rCNNs may be useful in artificial intelligence technologies, particularly those operating under resource constraints (e.g. [36, 54, 55]).

Reusing weights across time also reduces the passive costs of connections: In biological systems, connections need to be developed, accommodated in the body, and continually nourished, which requires energy and space, even when the network is idle. In artificial systems, similarly, there are costs of construction and space if neuromorphic hardware is used, and costs of memory storage if the network is emulated on a conventional computer. In both biological and articifical systems, the experiential data and energy required for learning a large number of parameters constitute additional costs. The need for large amounts data, energy, and time for learning, in fact, is among the most significant drawbacks of current neural network models. Recurrent models offer an avenue for limiting the number of parameters without limiting the computational depth or total computational budget for an inference.

### Learned lateral recurrent connectivity

As part of an exploratory analysis of the lateral connectivity in the BL networks, we observed that these models may learn recurrent connectivity profiles that resemble those in biological vision (see S1 Text). We found connectivity that could be interpreted as evidence for centre-surround computations and could support properties such as sparse representations [56], border ownership [50], contour integration [51], and end-stopping [57]. These analyses of recurrent connectivity offer a promising starting point for understanding recurrent computations in artificial visual systems and should be followed up by a detailed analysis of activity patterns in the models.

The observed lateral connections in our networks trained for object recognition also show a resemblance to the lateral connections of networks trained for contour integration tasks [30]. Given the different nature of these tasks, the similarity in lateral connectivity is surprising. This leads to the interesting hypothesis that there might be a subset of lateral computations that are useful across a range of visual tasks, at least in low-level visual areas. This would be consistent with the fact that a large range of objectives can be optimised to obtain simple-cell like features as observed in low-level visual areas. Such objectives include image classification performance [58], predictive coding [59], temporal stability [60, 61], and sparsity [56].

### Future directions

This study adds to a growing body of research on rCNNs as models of object recognition [25–29, 31, 32]. The rCNN model class could provide a unified basis for predicting stimulus-specific distributions of errors and reaction times in different sensory modalities and perceptual tasks, complementing previous work on recurrent processing in the decision-making literature.

Recurrent processing in human decision-making is often interpreted as serving to accumulate evidence. When the evidence consists in a noisy signal that reflects some variable of interest, the optimal inference procedure is to sum [33]. In dynamic real-world situations, however, the content of the signal varies over time, for example when the observer moves and previously hidden elements of the scene come into view. Perception is an ongoing inference process where a dynamic sensory stream meets a dynamic internal representation of the scene. Even for a static sensory input (as in the present study), each step of inference might depend on preceding steps, with sudden insights changing the course of the process. Recurrent neural network models can capture such processes and will be essential for understanding the recurrent computations of biological vision.

## Methods

### Deep neural network implementation

#### Architecture descriptions

All deep neural networks in these experiments were implemented using TensorFlow [62]. The baseline feedforward model (B), the recurrent model BL and the feedforward models parameter-matched to BL (B-K, B-F, B-D) are specified in detail in Table 3.

**Table 3.** Specification of network architectures

| Model | B | B-K | B-F | B-D | BL |
| --- | --- | --- | --- | --- | --- |
| Block 1 | F = 96, K = 7 | F = 96, K = 11 | F = 192, K = 7 | F = 96, K = 7<br>F = 96, K = 7 | (F = 96, K = 7) × 2 |
| Pool 1 | 2 × 2 max pooling | | | | |
| Block 2 | F = 128, K = 5 | F = 128, K = 7 | F = 256, K = 5 | F = 128, K = 5<br>F = 128, K = 5 | (F = 128, K = 5) × 2 |
| Pool 2 | 2 × 2 max pooling | | | | |
| Block 3 | F = 192, K = 3 | F = 192, K = 5 | F = 384, K = 3 | F = 192, K = 3<br>F = 192, K = 3 | (F = 192, K = 3) × 2 |
| Pool 3 | 2 × 2 max pooling | | | | |
| Block 4 | F = 256, K = 3 | F = 256, K = 5 | F = 512, K = 3 | F = 256, K = 3<br>F = 256, K = 3 | (F = 256, K = 3) × 2 |
| Pool 4 | 2 × 2 max pooling | | | | |
| Block 5 | F = 512, K = 3 | F = 512, K = 5 | F = 1024, K = 3 | F = 512, K = 3<br>F = 512, K = 3 | (F = 512, K = 3) × 2 |
| Pool 5 | 2 × 2 max pooling | | | | |
| Block 6 | F = 1024, K = 3 | F = 1024, K = 5 | F = 2048, K = 3 | F = 1024, K = 3<br>F = 1024, K = 3 | (F = 1024, K = 3) × 2 |
| Pool 6 | 2 × 2 max pooling | | | | |
| Block 7 | F = 2048, K = 1 | F = 2048, K = 3 | F = 4096, K = 1 | F = 2048, K = 1<br>F = 2048, K = 1 | (F = 2048, K = 1) × 2 |
| Readout | global average pooling<br>565 or 1000 category readout |  |  |  |  |
| Parameters | **11.0 million** | **39.8 million** | **40.0 million** | **28.9 million** | **28.9 million** |

Each row in the table represents a convolutional layer. F specifies the number of feature maps in the layer and K represents the height and width dimensions of the convolutional kernel. For BL, “(...) × 2” indicates that the same size convolutional kernel is applied twice, once to the bottom-up input (from the layer below) and once to the lateral input (from the same layer). All convolutions are applied with $1 \times 1$ stride and all max pooling is applied with $2 \times 2$ stride. The number of parameters are calculated for ImageNet models, ecoset models have slightly fewer parameters for the readout due to the smaller number of categories in ecoset.

The recurrent network (BL) is unrolled across time (Fig. 8) for eight time steps. At each time point in BL, the network receives an input image at the first layer and a readout is take from the last layer.

**Fig 8.**
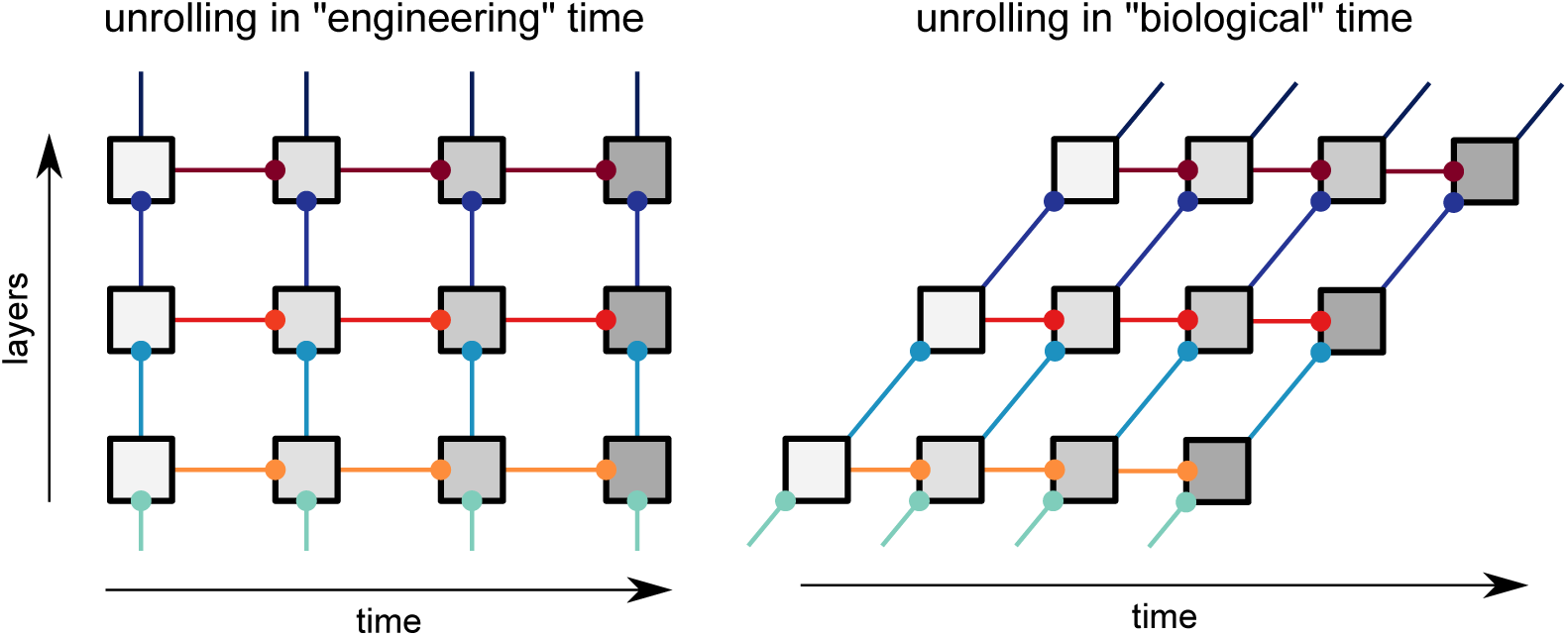
Network unrolling through time. Unrolling is shown for engineering time (left) and biological time (right). Each box represents a layer and the shading corresponds to its label in engineering time. Connections with the same colour represent shared parameters.

An additional feedforward model (B-U) was also trained. This model is identical to a BL network unrolled across time (for eight time steps) but, instead of sharing parameters across time, each convolution has unique parameters. Similar to BL, B-U has multiple input and output layers directly mapping to the input and output layers of BL at each time step. B-U has a total of 212.7 million parameters.

#### Unrolling recurrent networks across time

Artificial recurrent neural networks are typically implemented with feedforward connections taking no time and recurrent connections taking a single time step, we refer to this as *“engineering” time*. In comparison, all connections in biological neural networks should incur some time delay. A more biologically realistic implementation of a recurrent network may have every form of connection taking a single time step [25, 29]. We refer to this as *“biological” time*. Network unrolling in engineering time and biological time yield distinct computational graphs in the presence of top-down connections. However, for BL networks (which have lateral, but not top-down connections), unrolling in engineering time and biological time produce equivalent computational graphs (Fig. 8). Note that we neglect (1) computations that occur prior to the first feedforward sweep and (2) computations that cannot reach the readout before the final time step is reached. Based on the equivalent computational graphs for BL networks, we chose to use “engineering” time for the recurrent networks here and defined time as the number of complete feedforward sweeps that have occurred.

Note that in the unrolling scheme for BL (Fig. 8), each layer receives a time-varying feedforward input. This means that feedforward and recurrent processing happen in parallel. Alternatively, an rCNN could be unrolled such that all recurrent computations are performed within a layer and only the final output is passed to subsequent layers (e.g. [31]), resulting in recurrent and feedforward processing occurring in sequence. This implementation suggests that the onset of responses at later stages will be delayed when recurrence is engaged in earlier layers. However, experimental observations suggest that response onset is not delayed in later stages of the ventral visual pathway when recurrent processing is being utilised [24, 25]. These experimental findings motivate our unrolling scheme for BL, with recurrent and feedforward processing occurring in parallel.

#### Convolutional layers

We define the output from a standard feedforward convolutional layer at layer *n* on time step *t* as

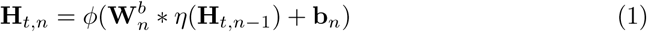

Where 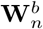 are the bottom-up convolutional weights for the layer and **b**_*n*_ are the biases. The convolution operation is represented as *. Optional max-pooling on the bottom-up input is represented by *η*. All other operations applied after the convolution are represented by the function *ϕ*. These operations include batch-normalisation [63] and rectified linear units in that order.

For a recurrent BL layer, the output is defined as

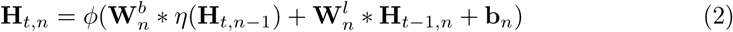

Where 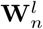 are the lateral recurrent weights.

For the recurrent networks, batch-normalisation is applied independently across time. Whilst this means that the networks are not truly recurrent due to unique normalisation parameters at each time step, this does not affect arguments related to parametric efficiency, as the numbers of parameters added by batch-normalisation at each time-step are negligible compared to the overall scale of the network. Approximately, 60,000 parameters are added across time due to batch-normalisation compared to 28.9 million parameters for the network as a whole.

In addition, we tested whether the use of independent batch-normalisation across time confers an additional performance advantage to recurrent networks by training B-D and BL on ImageNet without batch-normalisation. In this case, networks were trained using the same procedure but for only 25 epochs to prevent overfitting (as the removal of batch-normalisation reduces stochasticity in training). B-D and BL achieved a top-1 validation accuracy of 52.5% and 58.6%, respectively. This suggests that independent batch-normalisation across time does not explain the performance difference between feedforward and recurrent networks and even has a more beneficial effect for feedforward networks than recurrent networks (approximately 10 percentage point increase for B-D compared to a 6 percentage point increase for BL).

#### Network training

Before passing the images to the network, a number of pre-processing steps were applied. First, a crop was taken from the image, which was resized to 128 *×* 128 pixels. During testing and validation, a centre crop was taken from the image. During training, a random crop was taken covering at least one third of the image area. Further data augmentation was also applied in training, this included random left-right flips, and small distortions to the brightness, saturation and contrast of the image. Finally, the pixel values in the image were scaled from the range [0, 1] to be in the range [-1, 1].

B, BL and parameter-matched controls (B-K, B-F and B-D) were trained for a total of 90 epochs with a batch size of 100. B-U was trained using the same procedure but with a batch size of 64 due to its substantially larger number of parameters.

The cross-entropy between the softmax of the network category readout and the labels was used as the training loss. For networks with multiple readouts (BL and B-U), we calculate the cross-entropy at each readout and average this across readouts. Adam [64] was used for optimisation with a learning rate of 0.005 and epsilon parameter 0.1. L2-regularisation was applied throughout training with a coefficient of 10^*−*6^.

The code for models and weights for pre-trained networks are made available at github.com/cjspoerer/rcnn-sat.

### Defining accuracy in recurrent networks

As recurrent networks are unrolled across time, they have readouts at multiple time steps. This means that we must map from many readouts for a single image to one prediction. This leads to some ambiguity about how to produce predictions from recurrent networks for object recognition. Therefore, we conducted initial analyses to determine how to generate predictions from recurrent networks in the experiments described here.

One decision is how to select the time step to readout from the network, which we refer to as the network’s reaction time. A fixed time step could be chosen. For example, the readout could always be taken at the final time step that the recurrent model runs until. We refer to this as time-based accuracy.

Alternatively, we could select the readout to use based on when the model reaches some threshold. For example, the prediction is taken from the network once a certain level of confidence is reached. This confidence level could be defined by the entropy of the readout distribution where a lower entropy corresponds to a higher confidence. If the required confidence level is never reached then the final time step is selected as the reaction time. This is referred to as threshold-based accuracy. It should be noted that threshold-based accuracy can be implemented in recurrent networks using dynamic computational graphs that only execute up to the desired threshold. However, for our analyses we simply measure the time that it takes for the network to achieve a given level of entropy.

Once the decision time has been selected, we need to decide how to reduce the readout distribution across time. One method is to generate the prediction based solely on the readout at the network reaction time. We refer to this as the instantaneous readout. A second method is to generate the prediction from the cumulative readout up to the decision time, allowing the network’s predictions to be explicitly aggregated across time.

These different methods were compared using held-out data (Fig. 9). For ecoset the held-out data corresponds to the test set and for ImageNet this corresponds to the validation set, as the test set is not publicly available.

**Fig 9.**
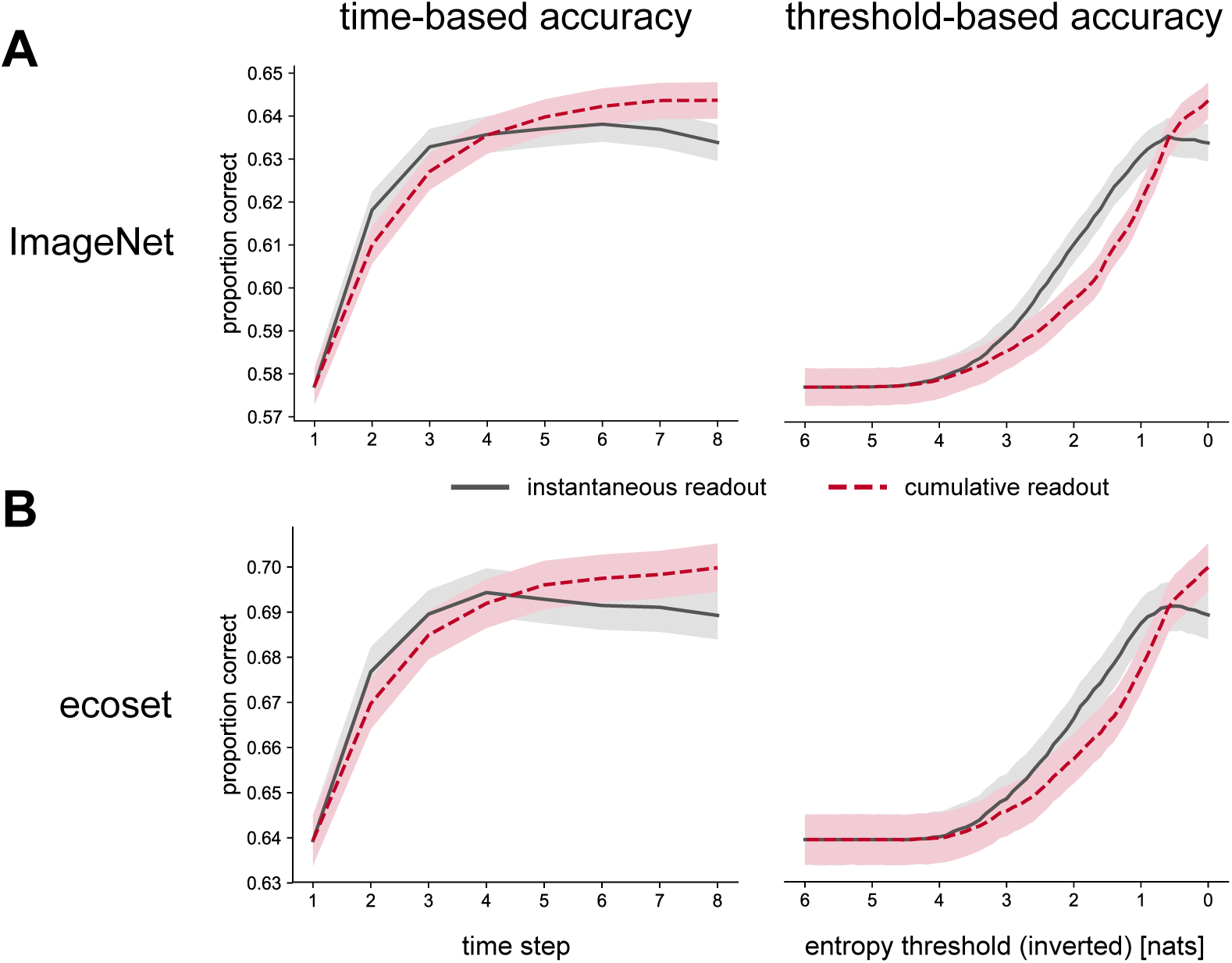
Task performance using varied definitions of predictions for recurrent models. Accuracies are given for models trained on (A) ImageNet and (B) ecoset using both time-based (left) and threshold-based (right) methods. Accuracies obtained from instantaneous readouts are shown with solid lines and results from cumulative readouts are shown with dashed lines. Shaded areas represent 95% confidence intervals obtained through bootstrap resampling.

For time-based methods, we see that the accuracy of the readout tends to increase across time. However, there is some drop-off in performance at later time steps if the instantaneous readout is used. One explanation for this pattern is that, by training the network to produce a readout at each time step, the network is encouraged to produce accurate predictions more quickly at the cost of higher accuracy at later time steps.

If a cumulative readout is used then accuracy improves more steadily across time, which is consistent with the smoothing effects expected from a cumulative readout.

However, cumulative readouts produce a higher overall level of accuracy than instantaneous readouts. This suggests there is some benefit of accumulating evidence across time for the performance of the network, even though the predictions themselves are not independent across time.

Similar results are seen when threshold-based accuracies are used. This reflects the fact that decreasing the entropy threshold will naturally lead to later time steps being increasingly utilised. Threshold-based accuracies also show a decrease in accuracy for instantaneous readouts at the lowest entropy levels. This is again due to worse performance at later time steps but also highlights an assumption of threshold-based accuracies that letting the network run for longer, to obtain higher confidence levels, will generate better predictions.

As a result of these analyses, all reported accuracies for recurrent networks refer to predictions based on cumulative readouts as these tend to produce the best performance.

### Behavioural experiments

#### Participants

Twenty healthy participants (16 female) aged 22-35 years (mean 26.62 years ±4.21) were recruited from the Medical Research Council – Cognition and Brain Sciences Unit volunteer panel. All participants had normal or corrected-to-normal vision, and reported no history of neurological or psychiatric disorders. The experimental procedure was conducted in accordance with the Cambridge Psychology Research Ethics Committee. Participants provided written informed consent and were compensated financially for participation.

#### Materials

We used the experimental stimuli from (Kriegeskorte et al. [65]). The stimuli presented to our participants were 96 colour photographs (250 *×* 250 pixels) of isolated real-world objects on a grey background. The objects included natural and artificial inanimate objects as well as faces and bodies of humans and nonhuman animals. Forty-eight pictures out of the 96 were animate objects, 12 human bodies, 12 animal bodies, 12 human faces and 12 animal faces. Twenty-four pictures out of the 48 inanimate objects were depicting man-made objects while the remaining 24 depicted natural objects.

#### Experimental procedure

The experiments were programmed using the Psychophysical Toolbox [66, 67] in Matlab (MathWorks, Natwick Inc) on a Dell Desktop PC computer. The participants were instructed to categorise “as quickly and as accurately as possible” objects according to the animate vs. inanimate categorical dichotomy. For each stimulus presentation, the participant had to press one of two keyboard keys as quickly as possible to indicate from which one of the two categories the stimulus was drawn. Each stimulus was presented exactly 6 times. Within the task, the order of the stimulus presentation was pseudo-random controlling for potential confounds related to stimulus presentation order. The trial onset asynchrony was 2 seconds and the stimuli were shown for a duration of 500 ms, providing the participant with 2s (including stimulus duration) to indicate the object’s category before the next object was presented.

### Fitting network reaction times to human reaction times

A cross-validated procedure was used to fit network models to human reaction times in the animacy discrimination task (as described in *Behavioural experiments*). The network models tested included B-D (ImageNet-trained and ecoset-trained), B-U (ImageNet-trained) and BL (ImageNet-trained, ecoset-trained and randomly initialised). A range of networks pre-trained on ImageNet that are popular in the engineering literature were also included [42–47]. The procedure involved two key steps, training the animacy discrimination readout and fitting the entropy threshold.

#### Training the animacy discrimination readout

To explain the human reaction times, animacy discrimination readouts were trained at eight points throughout the networks. The position of the first readout to reach a specified entropy threshold was taken as the network reaction time. For networks with multiple readouts (B-U and BL) readouts were trained in the same position as the original readouts. For feedforward networks without multiple readouts (B-D and pre-trained ImageNet models), a set of eight readouts were placed in an ordered sequence so that a similar number of additional computations were performed between any pair of adjacent readouts. Only a subset of layers were considered as candidate readout layers for the feedforward models trained without multiple readouts (Table 4 summarises the layers considered for each model).

**Table 4.** Subset of layers considered for training animacy discrimination readouts in single-readout feedforward models

| Model | layers considered for animacy readouts |
| --- | --- |
| B-D | ReLU layers |
| Inception-ResNet v2 | ReLU layers in the network stem, output of mixed concat layers, output of ResNet blocks, final ReLU layer |
| Xception | ReLU and add layers |
| NASNet | concat layers |
| DesnseNet-201 | concat layers |
| ResNet-50 | ReLU layers |
| VGG16 | ReLU layers |

To train the animacy readout, activations for each of the eight selected readout layers were taken in response to 899 training images (406 animate and 493 inanimate). These images were taken from a stimulus set of 1024 cropped images on a mid-grey background [68]. Images that also appeared in the behavioural experiment, or did not clearly depict animate or inanimate objects were removed from the training set. The remaining images were labelled as animate or inanimate.

The extracted activations underwent a step of dimensionality reduction, using principal components analysis (PCA), fitted on the training set, to project the activations into a 512-dimensional space. For recurrent networks, PCA was fitted for all time steps simultaneously. This simplified training the animacy readout as it reduces the number of parameters to be optimised. It also has the benefit that all network layers are reduced to the same dimensionality. Therefore, changes in the readout across layers cannot be explained by changes in the dimensionality of the input or (as a consequence) the number of the parameters in the readout.

A sigmoid animacy discrimination readout is then trained to maximise performance using activations for the training images projected in 512 dimensions. For the recurrent networks a recurrent sigmoid readout is trained across all time steps. The output of the recurrent readout at time step *t* ∈ {1..8} is defined as

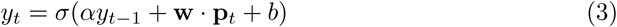

Where **p**_*t*_ are the loadings on the principal components at each time step, *α* is a recurrent parameter that allows evidence to be accumulated across time, **w** are the weights for the linear readout, *b* is the bias and *σ* is the sigmoid non-linearity. The initial readout state *y*_0_ was defined to neutral, such that *y*_0_ = 0.5. For feedforward networks, there is no parameter sharing across the layers, therefore, a separate sigmoid readout is trained for each readout layer.

The readout was optimised using batch gradient descent with Adam. The learning rate was set to 0.001 and the readout was trained for 1000 iterations. The loss was weighted for each class to account for the imbalance of classes in the training set.

This procedure was repeated 10 times, initialising the PCA and readout from different random seeds (note that a randomised method for PCA is used given the size of the original activation space [69]). For each random seed the PCA and animacy readout were used to produce responses to each of the 96 images used in the behavioural experiments, saving the results for each random seed.

#### Cross-validated procedure for entropy threshold selection

Entropy thresholds were used to extract reaction times for each of the 96 images used in the behavioural experiments. A double leave-one-out cross-validation procedure was used for fitting the entropy threshold. In each fold of the cross-validation procedure a single image (across all subjects) and subject (across all images) were removed as the test image and subject, respectively. The remaining 95 images across 19 subjects were taken as the training set.

The entropy threshold was found that maximised the correlation between network reaction times (averaged across random seeds) and human reaction times (averaged across participants) on the training set. Using the entropy threshold fitted on the training data, a predicted reaction time was extracted for the left out image and subject. The predicted reaction time was recorded for later analysis. This procedure was repeated until all subjects had a predicted reaction time for every image, fitted using independent data.

The cross-validated network reaction times were then compared to human reaction times for each subject individually using Pearson correlation. Pearson correlation was used as we expect the relationship between human and network reaction times to be linear. The correlation coefficient across human subjects was averaged and a paired permutation test (with 10,000 permutations) was used to test for significant differences in the mean.

### Extracting lateral-weight components

We analyse the lateral connectivity of the network by decomposing the lateral weights in the network into lateral-weight components. To do this, we focus of the 7 *×* 7 weight templates that connect each of the feature maps within the first layer of the network. There are 96^2^ weight templates in total connecting every feature map to each other in both directions (including self-connections from a feature map to itself). We focus on the first layer of the network as the corresponding bottom-up weights are easier to interpret and recurrence is arguably best understood in early regions of the visual system (corresponding to early layers of the network).

Firstly, the weight templates are normalised such that the vector of the flattened weight template has unit length. After normalisation, the lateral weights are processed using principal components analysis (PCA) where each weight template is considered as an individual sample. The first five components resulting from the PCA are used as the lateral-weight components for the analysis.

## Supporting information

**S1 Fig. Median accuracy in the human reaction times task**. The median accuracy (median across participants proportion correct response in classifying animate vs. inanimate) was 1 for 91 out of the 96 stimuli.

**S1 Text. Further explorations of learned lateral connectivity in rCNNs**.

