## Supporting Information for "Recurrent neural networks can explain flexible trading of speed and accuracy in biological vision"

### S1 Fig: Accuracy in the human reaction times task

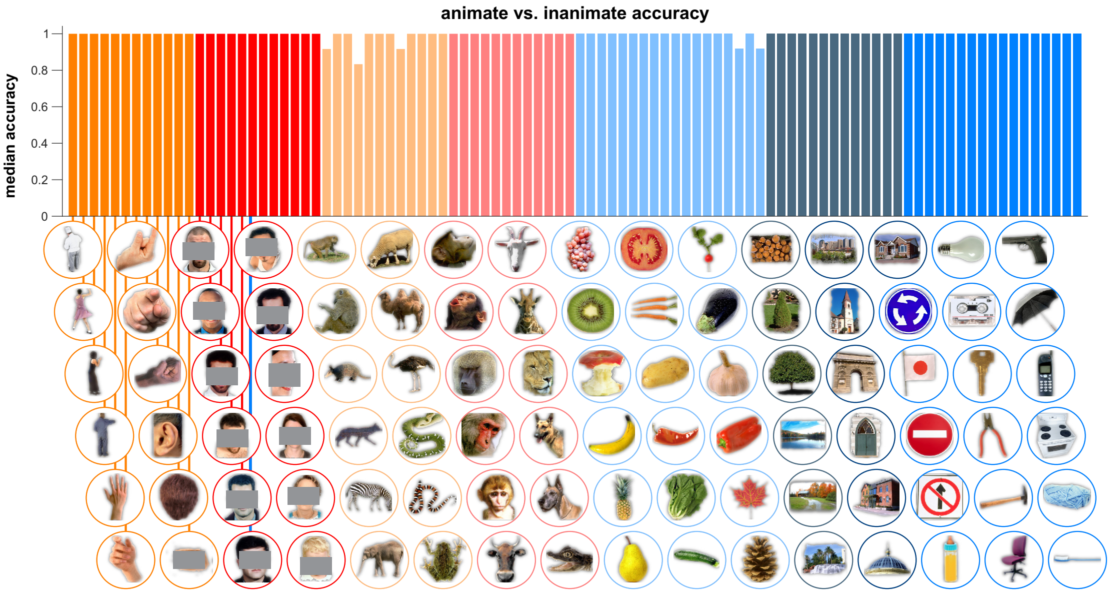


**S1 Fig. Median accuracy in the human reaction times task.** The median accuracy (median across participants proportion correct response in classifying animate vs. inanimate) was 1 for 91 out of the 96 stimuli.

### S1 Text: Further explorations of learned lateral connectivity in rCNNs

#### Local inhibition/excitation

The lateral-weight component explaining the most variance in the network corresponds to local inhibition and excitation. Near inhibitory connections could be used to generate sparse representations, similar to visual cortex [1].

To further understand how inhibitory connectivity relates to the properties of bottom-up features, we correlated the bottom-up weight templates of features connected by lateral-weights with strong negative loadings on the first component (defined as the lowest percentile of loadings on the component). We found a median correlation of -0.16 between bottom-up features with local inhibitory recurrent connections. This value significantly differed from zero (Wilcoxon signed-rank test, $p < 0.001$), suggesting that dissimilar features inhibit each other in the network, possibly increasing the sparsity of the representation.

### Centre-surround antagonism

Centre-surround antagonism is a well-studied feature of biological vision and is most often seen in the context of near excitation and far inhibition. In these arrangements, a unit will be excited if a preferred stimulus is detected in the centre and suppressed if the preferred stimulus appears in the surround.

In the lateral-weights of the network, we see centre-surround antagonism in both the classical arrangement of near excitation and far inhibition and the non-classical arrangement of near inhibition and far excitation (Fig. 6, component 3). However, features connected with non-classical centre-surround connectivity (highest percentile of loadings on component 3) had a median negative correlation of -0.04, which significantly differed from zero (Wilcoxon signed-rank test, $p = 0.003$). Non-classical centre-surround connectivity in the network, thus, could still lead to reduced responses if a preferred stimulus is detected in the surround, like classic centre-surround connectivity, but due to reduced excitation rather than increased inhibition.

### Cardinal antagonism

Vertical and horizontal antagonism are also observed in the network (Fig. 6, component 2 and component 4). We collectively refer to vertical and horizontal antagonistic weight templates as cardinal antagonism. This type of interaction leads to excitation if a feature is detected to one side of a unit and leads to inhibition if that same feature is detected on the opposite side. This type of asymmetry could be useful for developing border ownership cells [2], which have varying levels of response, depending on which side of an edge corresponds to an object or background surface.

A unit that detects an edge between two surfaces could show properties of border ownership if it receives recurrent input carrying information about the spatial extent of the two surfaces meeting at the edge. We see examples of this type of connectivity in the network. For instance, feature 76 is sensitive to purple-green edges and it receives input from feature 78, which prefers diffuse purple features (Fig. 6, component 4). The recurrent connectivity between them is cardinally antagonistic such that the unit detecting the purple-green edge is only excited if a diffuse purple feature is detected on the purple side of the edge.

### Perpendicular antagonism

Perpendicular antagonism is observed in this network where there are excitatory recurrent connections along one orientation and inhibitory recurrent connections along the orthogonal orientation (in both directions). This type of connectivity is consistent with association fields that could support contour integration [3].

Studying the feature maps that most heavily load on these components, we find that feature maps that detect gradients in similar orientations with edges in phase have collinear inhibition and orthogonal excitation (Fig. 6, component 5). In comparison, we see collinear excitation and orthogonal inhibition when feature maps are detecting gradients that have similar orientations but opposite phases.

Collinear excitation may be expected between features detecting gradients in similar directions because the presence of such features is consistent with a continuous contour. However, collinear inhibition is consistent with end-stopping behaviour observed in complex cells of visual cortex [4]. In this case, cells were observed that have suppressed firing rates if edges extend beyond the classical receptive field of the cell.
